# Role of Speed Regulation and Speed Modulation in Velocity-Field Based Control

**DOI:** 10.1101/2025.09.15.676406

**Authors:** Rezvan Nasiri, Lyndon Tang, Michael Goldfarb, Arash Arami

## Abstract

The velocity vector field (flow) controller is a well-established control strategy for lower limb exoskeletons. In this paper, we analyze this controller and propose modifications to improve its performance. We demonstrate that flow control acts as a variable proportional-derivative error regulator, where the parameter Γ represents the desired norm of the hip-knee joint velocity vector (path speed). Based on this, we introduce two modifications to Γ: (1) a constant Γ set to the mean desired path speed, and (2) a variable Γ that mimics natural path speed during unassisted walking. We compared the modified flow controllers with a slow-Γ version in experiments involving seven participants walking on a treadmill at 0.6*m/s*, 0.8*m/s*, and 1.0*m/s*. Compared to the slow-Γ controller, the RMS tracking error decreased by 30.7*±* 11.3% and the range of motion of the knee increased by 48.2 *±* 5.5% for the mean-Γ controller, while the variable-Γ controller had 32.4 *±* 14.7% smaller RMS error and 50.5 *±* 6.5% larger range of motion of the knee. Additionally, the slow-Γ controller consistently applied resistive power, whereas participants reported more comfortable and natural gait with the modified controllers. We also compared them with the original tuning of flow controller, with results indicating superior performance from the proposed modifications. These findings demonstrate effectiveness across different walking speeds and offer a tuning strategy for future flow controller use.

## I. Introduction

Lower limb exoskeletons have shown great potential to improve gait rehabilitation in individuals with motor deficits due to stroke or neurological conditions [1], [2]. Despite their advantages (e.g., assisting with walking tasks during physical therapy to improve the rehabilitation outcome), there are still challenges in the design of controllers that prevent users from fully harnessing the maximum potential of the exoskeleton. Numerous approaches have been developed for controlling lower limb exoskeletons, including trajectory-based controllers [3] and force controllers [4]. However, time-based trajectory tracking has been shown to be ineffective for assisting individuals with partial motor impairments, as it fails to fully engage the users in contributing to their gait [1]. These controllers could not synchronize with the user’s intended motion and often fail to account for the level of needed assistance for each user. Event-based and gait-phase-based trajectory tracking [5], feedback control based on neuromuscular reflexes [6], and hybrid feedforward assistive torques [7] have demonstrated improved performance stem from enhanced synchronization with the user intent.

To improve robotic rehabilitation outcomes, another strategy is to encourage maximal user engagement in the control and execution of the movement, while limiting the exoskeleton assistance to only when it is needed [8], [9]. Accordingly, several existing assist-as-needed (AAN) controllers have been designed; examples include the path controllers [10], [11], the velocity vector field controller [12], controllers that adapt the exoskeleton trajectory in real-time [13], and controllers that work in energy space, like energy shaping control [14], [15], and virtual energy regulator [16], [17].

In 2009, Duschau-Wicke et al. [10] introduced an early AAN controller known as the path controller, which applies joint-level impedance towards the nearest joint configuration on a reference path when the deviation exceeds a defined channel threshold, allowing users to retain some gait variability and actively contribution to movement. Goldfarb et. al later implemented the path controller in their lower limb exoskeleton (i.e., Indego) for walking at 0.8*m/s* [11]. They subsequently developed Velocity Vector Field (or flow) controller and compared it with the path controller during overground walking at the same speed [12].

The flow controller is easy to implement and its experimental results show a promising performance. Accordingly, it was implemented in other studies such as knee exoskeleton [18], full body exoskeleton for stroke patients [12], and upper limb exoskeletons [19]. Jamšek et al. integrated the flow control with probabilistic movement primitives for an upper limb exoskeleton for reaching tasks [20]. The controller has been proven in many gait rehabilitation applications to be reliable and effective in providing assistive torques to the user. However, tuning of the controller parameters have not been effectively investigated. A task-dependent or personalize tuning may lead to better regulation of joint kinematics, and improved coordination with the user to apply assistance only as-needed. In this paper, we analyze the flow controller and propose a modification to improve its performance in lower limb exoskeletons during gait.

Specifically, we show that: (1) the flow control’s velocity regulation acts as a damping force, potentially hindering natural, and beneficial swing phase velocity changes, (2) we propose modifications to the Γ parameter of flow controller to enhance the velocity regulation, and (3) we experimentally compare flow controllers with different fixed and variable Γs on seven participants walking with an exoskeleton at three different treadmill speeds to demonstrate a method for systematic tuning of this assistive controller.

## II. Methods

In this section, we rearrange the original formulation of the flow controller [12], and show that it is a weighted sum of the kinematic error and its time derivative. Then, we propose a method for modification of the flow controller to improve its performance at different gait speeds.

### A. Flow controller

The velocity vector field controller (i.e., flow controller) presented in [12] applies torque (based on a flow field) to regulate the joint velocity (*ω*) to the reference velocity (*ω*_*ref*_) as Eq. 1A where positive scalar *C*_*d*_ is the drag coefficient, or controller gain. The reference velocity (as in Fig. 1) is a weighted sum of kinematic error (**e**) w.r.t. the minimum distance on the desired path and its corresponding normalized tangent vector (**t**) on the desired path (Eq.1B)

**Fig. 1.**
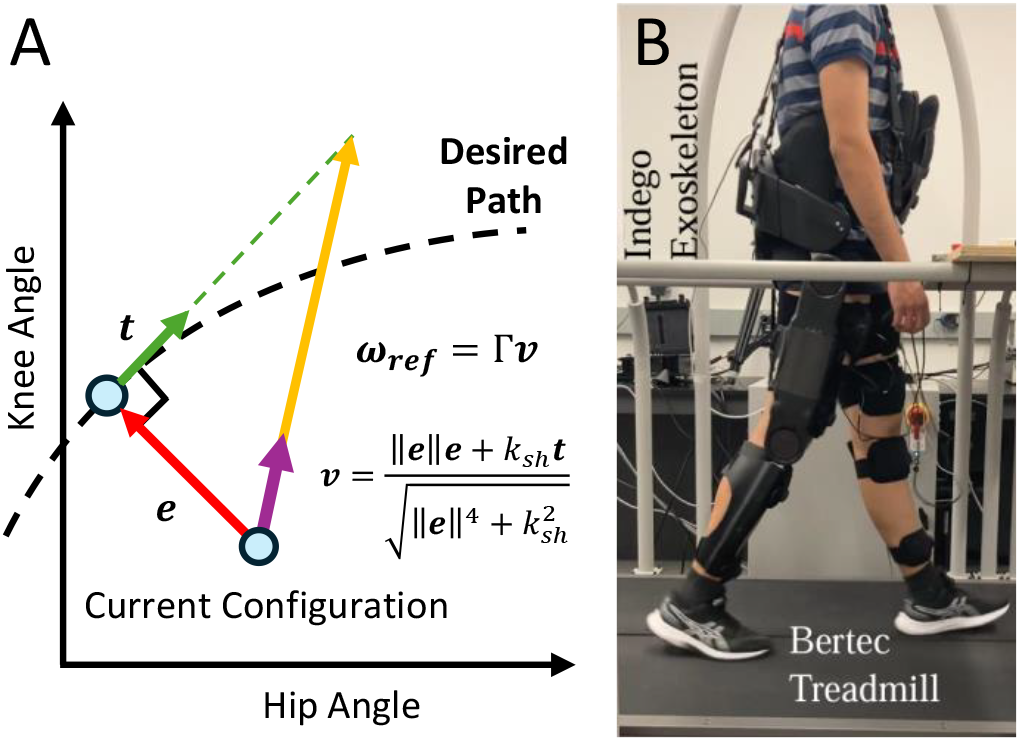
(A) The velocity vector field (flow controller) parameters; **e** is error vector, **t** is unit tangent vector to the path. (B) The experimental setup.

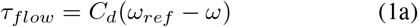

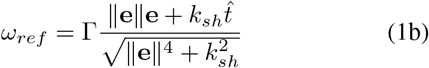

where Γ determines the magnitude of the velocity reference, and *k*_*sh*_ is the flow shaping gain. The controller is only active during the swing phase which is detected by a state machine.

### B. Mathematical analysis

Now, consider **q**_*ref*_ as the desired joint angles that define the path and its induced time derivative 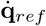. Considering the error vector as **e** = **q**_*ref*_ − **q** and replacing the tangent vector (which is the normalized desired velocity over the path) as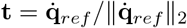, Eq.1 can be rewritten as

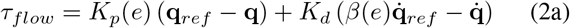

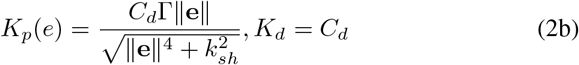

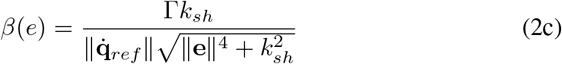

where we call *β*(**e**) speed modulation gain. This equation is a summation of two terms: (1) a variable positive gain multiplied to a configuration error plus (2) a positive gain multiplied to a velocity error. Eq. 2 indicates that flow controller functions as an impedance controller, regulating the behavior of the exoskeleton along the path at a modulated desired speed of 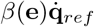. From this point of view, the flow controller is very similar to the path controller [10].

### C. Path speed modulation

Based on Eq.2, for configurations near to the desired path (i.e., **q**_*ref*_ ≊ **q** *⇒* **e** ≊ 0), the controller tries to regulate the gait by maintaining the second term; i.e., 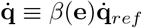. Applying the norm to both side of this equality, the control objective can be obtained as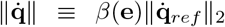. For **e** ≊ 0, the speed modulation gain is 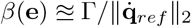, and therefore, the control objective can be rewritten as 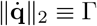. In other words, the controller regulates the norm of the joint angular velocity vector, referred to as the *path speed*, to Γ, which we call the *nominal path speed*. We hypothesize that, to achieve a natural and comfortable gait, Γ should be a function of the desired path speed (i.e., 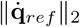). Therefore, we propose setting Γ in the flow controller (Eq.1) using one of two methods: (1) a constant Γ based on the average desired swing-phase path speed, or (2) a variable Γ that adapts to path speed variation in healthy gait.

Fig. 2A compares the average and the variation of path speed during swing phase of a normal gait at 0.8*m/s* with the suggested value for Γ at the experimental walking speed of 0.8*m/s* in [12]; the path speed reported in this figure is the average obtained across 50 gait cycles of a representative participant walking on treadmill without an exoskeleton at 0.8*m/s*. The plot shows that the average path speed during swing phase (Γ = 265*deg/s*) is much higher than the suggested value for Γ = 167*deg/s*.

**Fig. 2.**
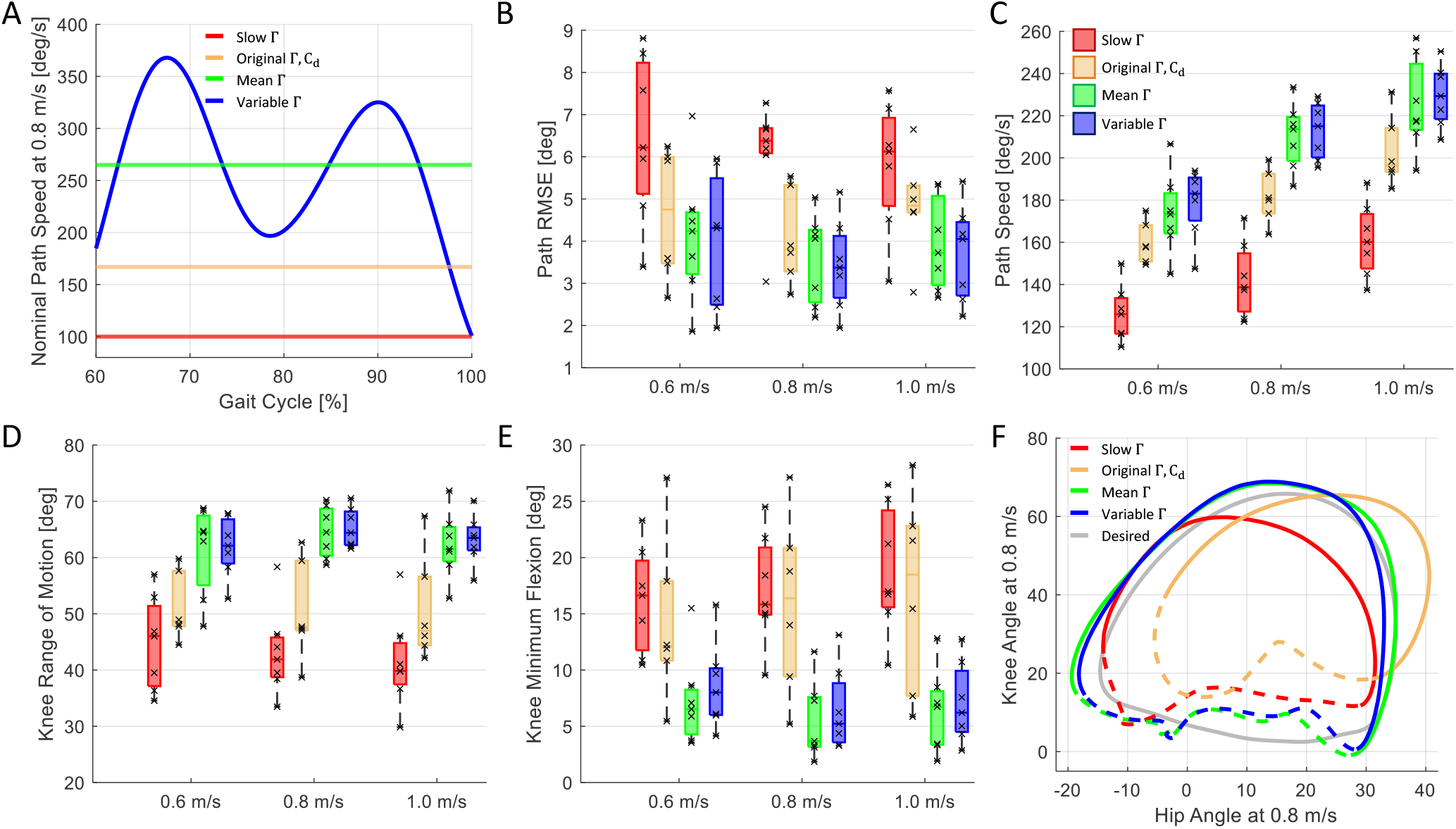
The experimental results: (A) The desired path speed during the swing phase at 0.8*m/s*. (B) The path RMS error. (C) The resultant path speed during the swing phase. (D) Knee minimum flexion angle. (E) The knee joint range of motion. (F) The resultant gait path for a representative participant at 0.8*m/s* during stance (dashed line) and swing (solid line); note that the gait path patterns are similar across different participants.

### D. Experimental setup, protocol, and validation

To investigate the effect of modulating the nominal path speed, Γ, we tested the flow controller with Γ = 100*deg/s* representing slow walking, Γ set as the mean observed path speed, and a variable Γ equal to the instantaneous path speed of walking without an exoskeleton, shown in Fig.2A.

Each controller was tested at three walking speeds, 0.6*m/s*, 0.8*m/s*, and 1.0*m/s*. To emphasize the effect of Γ, in the first experiment, we chose a drag coefficient of *C*_*d*_ = 0.065*Nms/deg*, five times larger than the suggested value in [12]. The other parameter, *k*_*sh*_ = 16 and implementations, including the state-machine to enable the controller during swing phase, and definition of the reference error as the minimum distance to the path were the same as [12].

Seven able-bodied participants (4 males and 3 females, age: 24.1 *±* 1.4 years, body mass: 76.6 *±* 12.6 kg, and height: 180.6 *±* 2.2 cm; mean±std) participated in the first experiment. The participants provided informed consent prior to the experiment. The study protocol was conducted in accordance with the Declaration of Helsinki and reviewed and approved by the University of Waterloo Research Ethics Board (ORE#41794).

In this experiment, we implemented the flow controller on an Indego Explorer exoskeleton (ekso Bionics, USA), with active hip and knee joints providing assistance in the sagittal plane. The joint angles and applied motor torques are measured at 200*Hz*.

The experiment consisted of a training session on day 1 and a test session on day 2. During the training session, participants were given at least 30 minutes to familiarize themselves with the exoskeleton and the controllers while walking on a treadmill (Bertec, USA) at speeds of 0.6*m/s*, 0.8*m/s*, and 1.0*m/s*. The test session consists of three consecutive trials walking on the treadmill for 4 minutes at above-mentioned speeds with a 30-second rest between the trails. Each trail started with (1) passive walking with zero exoskeleton motor torques, (2) slow Γ, followed by (3) fixed, averaged Γ, and ended with (4) variable Γ, each for 1 minute. For simplicity, the three control scenarios are referred to as the Slow, Mean, and Variable controllers.

Furthermore, we conducted a second experiment with six participants (3 males and 3 females, age: 22.8 *±* 2.3 years, body mass: 69 *±* 14.4 kg, and height: 175.6 *±* 7.34 cm). The participants followed the same protocol as in the first experiment, with the Slow controller replaced with the Original controller (i.e, with the original controller parameters, Γ = 167*deg/s, C*_*d*_ = 0.016*Nms/deg, k*_*sh*_ = 16). Additionally, the drag coefficient for the Mean and Variable controllers was also set to the original value.

Five metrics are used to evaluate the proposed modifications. The step-average root mean square error (RMSE) of configuration (i.e., the norm of difference between current joint angles and the nearest point on the reference path) and the average path speed assess how well the controller meets its regulation goals.We also evaluate minimum knee flexion and knee range of motion, two gait metrics relevant to rehabilitation. Minimum knee flexion is evaluated when the controller is active, and indicates whether the swing leg sufficiently extends before the stance phase [21]. Knee range of motion helps distinguish natural gait from shuffling or low-foot-clearance steps, which may increase the risk of fall.

Additionally, we assess the power supplied by the exoskeleton to quantify user-exoskeleton interaction. The sign of power indicates whether the exoskeleton is assisting (positive power) or resisting (negative power), i.e. absorbing energy. The power supplied at joint *k*, is computed as

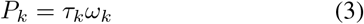

where *τ*_*k*_ is the motor torque, and *ω*_*k*_ is the joint angular velocity. While instantaneous power dissipation can reflect proper tracking assistance, consistent energy absorption may indicate undesired antagonistic behaviour. To summarize the net effect over a gait cycle duration (*T*), we compute the total energy supplied per step as

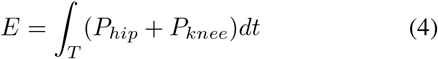

Group-level differences between the experimental results across the three controllers, were evaluated with *Friedman* test. Post-hoc *Wilcoxon Signed Rank* tests with a significance level of *α* = 0.05 and *Bonferroni* correction were done to analyze the statistical differences between the controllers. Differences in the metrics are reported as mean *±*SEM.

Comparisons between the results from the two experiments were made by first testing with a 9-group Kruskal Wallis test for group-level differences, followed by a *Wilcoxon Rank Sum* test with significance level of *α* = 0.05 and *Bonferroni* correction. Differences were reported between the participant averages for each experiment.

## III. Result and Discussion

Here, we first analyze the effects of the low, mean, and variable nominal path speeds of the flow controller (with high drag coefficient). Subsequently, we compare the effect of Γ across controllers with low *C*_*d*_ values, then compare the original controller settings with the ones mentioned above.

When comparing the metrics during the high-gain experiment, all five metrics passed the 9-group Friedman test with *p <* 0.002, indicating significant differences in the performances at the group level. In the second experiment, with a lower drag coefficient, the average path speed, range of motion, and minimum flexion angle of the knee, as well as the energy supplied passed the Friedman test with *p <* 0.007.

### A. Effect of Γ regulation in high C_d_ scenario

Both the Mean and Variable controllers led to reductions in the average path RMSE compared to the Slow controller, demonstrating better regulation of the joint configuration to the desired path, as shown in Fig.2B. The path RMSE decreased on average by 30.7 *±* 11.3% for the Mean Controller, and 32.4 *±* 14.7% for the Variable controller, relative to the path RMSE of the Slow controller. As expected, the average path speed during the swing phase increased significantly by 42.2 *±* 2.5% for the Mean controller to 205.5 *±* 6.5*deg/s* (*p* = 0.016), and by 45.7 *±* 3.4% for the Variable controller to 210.0 *±* 5.0*deg/s* (*p* = 0.016), when compared to the Slow controller (144.7 *±* 6.0*deg/s*), indicating that the Slow controller allowed greater path errors in order to regulate the path speed. There were no significant differences in the path RMS errors or the average path speeds between the Mean and Variable controllers.

The change in path error with different values of Γ is explained by the fact that, aside from acting as the nominal path speed, Γ also determines the magnitude of the reference joint velocity vector when the path error is large. As a result, in situations where the user is far from the path, the control torques also scale with Γ.

Fig. 3 shows the average path speed measured during the swing phase for different controllers at different gait speeds. The trajectories have similar amplitude patterns as the Variable Γ reference pattern in Fig. 2A. The Mean and Variable controllers both had significantly higher peak path speeds, and the path speed trajectories aligned better with the path speed observed during passive walking. As a result, the RMS difference in the path speed trajectories with respect to the average path speed during passive walking was significantly smaller by 45.3 *±* 4.8% for the Mean (*p* = 0.016) and 38.9 *±* 6.4% for the Variable controllers (*p* = 0.016) compared to the Slow controller.

**Fig. 3.**
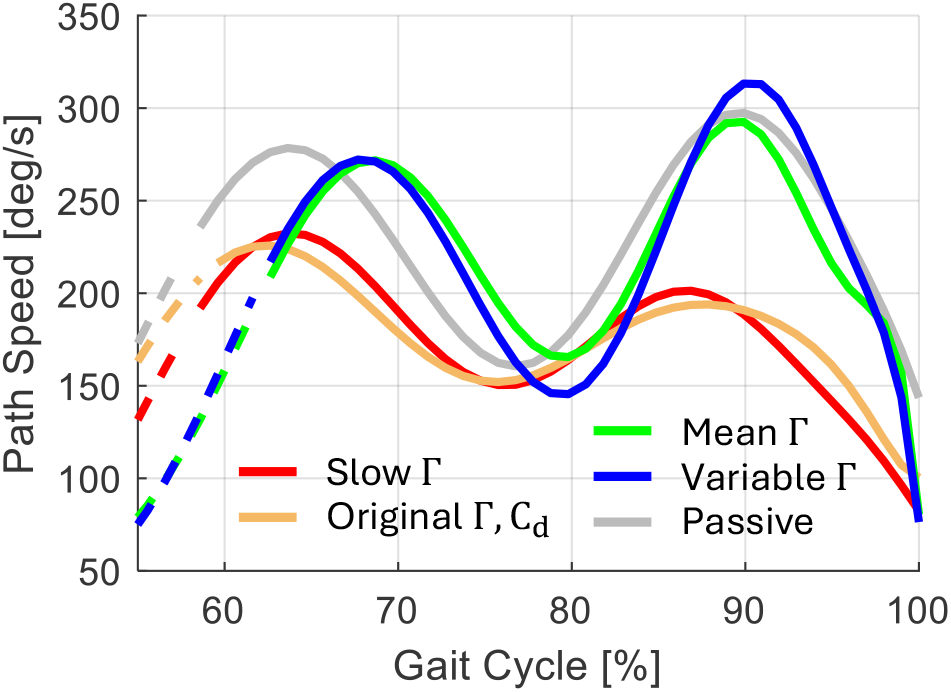
The average path speed trajectory of the Mean and Variable controllers has higher peak amplitudes than the slow controller, closer to the amplitude of the path speed during passive walking.

The significantly higher path speed of the Mean and Variable controllers compared to the Slow controller resulted in significant improvements in the gait kinematics. Participants had significant increases in their knee range of motion, shown in Fig. 2E, by 48.2 ± 5.5% for the Mean controller (*p* = 0.016), and by 50.5 *±* 6.5% for the Variable controller (*p* = 0.016). There were also significant decreases in the minimum flexion angle of the knee, shown in Fig. 2F, by 65.5 *±* 3.8 for the Mean controller (*p* = 0.016) and 59.3 *±* 4.2 for the Variable controller (*p* = 0.016), compared to the Slow controller. The Slow controller resulted in the smallest range of motion of all control scenarios of 42.9 *±* 2.90*deg* and a crouched gait, indicated by the minimum knee flexion angle of 17.6 *±* 1.9*deg*, far from full knee extension at 0*deg*.

Based on the participants feedback, the overall behavior of flow-controller with Γ = 100*deg/s* felt like a viscous damper, slowing them down, while the modified controllers felt more natural and comfortable. This indicates that the participants were trying to walk at a faster pace compared to the nominal path speed of the Slow controller, but the controller penalized the joint velocities and inhibited a natural gait with full range of motion and knee extension. The distortion of the knee flexion throughout the path cycle is shown for a representative participant in Fig. 2F with a reduced knee range of motion along the upper right section of the path, and incomplete knee extension at the heel strike in the bottom right of the cycle. The reduction in average path speed is also shown in Fig.3, where the path speeds of the slow and original controllers are much lower than the speeds for passive walking particularly during the swing phase.

The analysis of the power absorbed and supplied by the exoskeleton suggests the Slow controller’s negative impact on gait kinematics, from a lack of coordination between the user and controller. This controller constantly absorbed power, acting as a damper, as reported by participants. By convention, negative power and energy in this analysis indicates energy absorbed by the exoskeleton, while positive values indicate energy supplied.

As shown in Fig. 4A, the Slow controller produced negative power throughout swing phase for this representative participant, indicating continuous energy dissipation, resulting in an increase in user’s effort to maintain motion, consistent with the reported damping behaviour by the user. In contrast, the Mean and Variable controllers, which regulate higher path speeds, alternated between positive and negative power contributions, offering both assistance and resistance during swing.

**Fig. 4.**
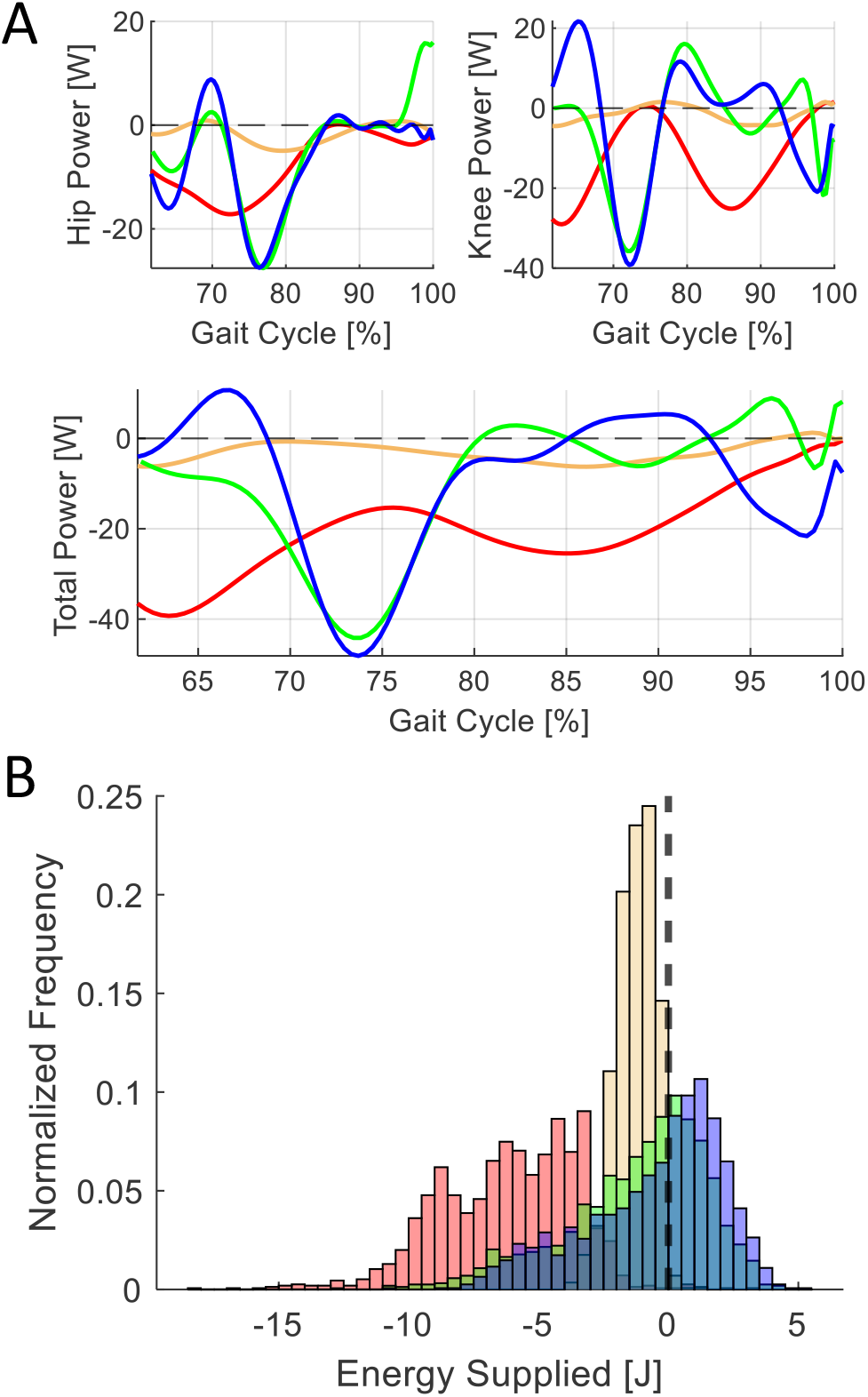
(A) The power supplied by the exoskeleton is negative throughout the swing phase of gait for the slow and original controllers, indicating that the exoskeleton is opposing the direction of motion. (B) The average energy supplied per step is significantly lower for all participants with the slow controller compared to the mean and variable controllers.

As a result, the average energy supplied by the exoskeleton was significantly more negative for the Slow controller (−6.16 *±* 0.56*J*) compared to the Mean (−1.21 *±* 0.77*J, p* = 0.016) and Variable (−0.50 *±* 0.81*J, p* = 0.016) controllers, and no significant difference between the Mean and Variable controllers. Fig. 4B shows the histograms of energy supplied per step for all participants at all walking speeds. The mode of the distributions of energy supplied per step were above zero for the Mean and Variable controllers, whereas, the mode of the distribution was negative for the Slow controller. All participants demonstrated this negative shift in the average energy supplied per step with the Slow controller, suggesting that all of the participants were walking faster than the controller was designed to regulate and expending energy to overcome the viscous damping effect.

### B. Effect of Γ regulation in low C_d_ scenario

In the second experiment with six participants and a low drag coefficient, trends in the metrics showed that higher values of Γ led to improvements in knee flexion and extension. Although the low sample size limits rigorous statistical analysis, a Friedman test revealed group level differences. Compared to the Original controller, the average path speed increased by 5.26 *±* 1.21% and 6.81 *±* 0.98% for the Mean and Variable controllers, respectively, with corresponding decreases in path RMSE by 3.15*±*4.82% and 8.07*±*5.46%. The range of motion of the knee also increased by 7.79 *±* 0.66% and 9.34 *±* 0.94% for the Mean and Variable controllers, respectively, while the minimum knee flexion angle decreased by 20.6 *±* 4.44% and 20.5 *±* 5.59%.

The energy supplied by the exoskeleton per step was more negative for the Original controller (−1.27 *±* 0.17*J*), which had the lowest Γ value, compared to the Mean (−0.332 *±* 0.210*J*) and Variable controllers (−0.289 *±* 0.208*J*), (*p* = 0.0312 for both). Both the Mean and Variable controllers had less negative values for the energy supplied with the low-gain version of the controller, indicating less resistance from the exoskeleton to the motions of the user.

Inspecting Eq. 2b shows that both the drag coefficient (*C*_*d*_), and Γ play important roles in scaling the control torque associated with the path error. Accordingly, path tracking error and knee kinematics improved in the first experiment, where *C*_*d*_ was set to five times the value used in the second experiment.

### C. Does drag coefficient affect Mean and Variable controllers?

A Kruskal Wallis test comparing the high-gain versions of each of the Mean and Variable controllers to their low-gain counterpart revealed significant group-level differences in average path speed, the RMS difference in path speed trajectories relative to passive walking, and knee range of motion (*p <* 0.05). Compared to the Original controller, the Mean and Variable controllers exhibited increases in average path speed of 6.30% and 7.08%, respectively, and reductions in path RMSE of 15.9% and 24.4%. Additionally, knee range of motion increased by 14.5% for both controllers, and the minimum knee flexion decreased by 51.5% and 44.2% for the Mean and Variable controllers, respectively. Although post-hoc rank sum tests suggested improvements with higher drag coefficient, these differences did not reach statistical significance. While this might be interpreted as tuning *C*_*d*_ is less importance than tuning nominal path speed (Γ), we refrain from making such a claim. It is important to clarify that our experiments were primarily designed to demonstrate the feasibility and effectiveness of the proposed Γ-tuning for the flow controller, rather than to establish statistical significance for the effect of individual controller parameters. Both the the nominal path speed and the drag coefficient can increase the magnitude of control torque, however, they modulate the torques differently. Γ, which is the focus of this study, can scale and rotate the exoskeleton torque vector by scaling ([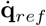 in Eq. 2), thereby altering the ratio of torques applied to hip and knee. In contrast, *C*_*d*_ only scales the control torque vector without affecting its direction.

### D. Comparison of high-gain Mean and Variable controllers with the original flow controller

Here, the experimental results of the Original controller (*C*_*d*_ = 0.016*Nms/deg* and Γ = 167*deg/s*) in the second experiment were compared to the Slow, Mean, and Variable controllers of the first experiment. A Kruskal Wallis test with a significance level of *α* = 0.05showed group level differences for all 5 metrics (*p <* 0.05).

The Rank Sum tests show that the effects of nominal path speed on the controller align with trends observed in the previous experiment: as nominal path speed increased, mean knee range of motion increased, and minimum flexion angle decreased. Compared to the Original controller, the Slow controller showed a significantly lower average path speed by 21.1% (*p* = 0.0023), while the Variable controller had a 14.4% higher path speed (*p* = 0.0082). RMS deviation of the path speed during passive walking was significantly larger for the Slow (167%, *p* = 0.0012) and Variable controllers (57.6%, *p <* 0.005), compared to the Original controller. Path RMSE was 34.5% higher for the Slow controller (*p* = 0.0513), but reduced by by 13.9% and 18.6% for the Mean, and Variable controllers (with higher Γ), respectively. The Variable controller increased knee range of motion by 25% (*p* = 0.014), while the Mean controller reduced the minimum knee flexion angle by 60% (*p* = 0.014), both compared to the Original controller.

The energy supplied per step by the exoskeleton also establishes a trend with the nominal path speed; the net energy supplied increases with increasing nominal path speed. The energy supplied by the Original controller was 385% larger compared to the Slow controller (*p <* 0.002). In Fig. Fig. 4A, the instantaneous power trajectory of the Original controller was negative throughout the swing phase of the gait cycle for the representative participant, showing that, like the Slow controller, the Original controller behaved as a viscous damper and dissipated energy from the user, suggesting that the nominal path speed was too low for the participant’s intended walking speed, and can be increased to improve the knee range of motion and prevent crouch gait.

### E. Tuning the nominal path speed

The Mean and Variable Γ controllers outperformed the Original and Slow flow controllers, offering new avenues for tuning the flow controller. Despite their improved performance over the original tuning, two undesirable behaviors were observed in some participants. First, a mild foot-stomping-like behavior was seen at higher speeds (0.8*m/s* and 1.0*m/s*), particularly when walking with shorter step lengths. For participants #2 and #3 in the first experiment, this resulted in increase in vertical ground reaction force (GRF) of 3.4% and 2.1% of body weight, respectively, during early stance (approximately 10% − 25% of gait cycle) when using the Mean controller, compared to the passive condition (i.e., walking with exoskeleton with zero torque). Second, we observed an excessive knee extension in some participants during the late swing when walking at 0.8 and 1*m/s*. For instance, for participants #2 and #3, minimum knee flexion reached an average of 0.02*deg* and 3.16*deg* respectively, when using the Mean controller.

To address these issues, we tuned the Mean controller by applying a corrective gain of 0.8 to the average path speed and conducted an experiment on two selected participants (#2 and #3) to assess the effect of this correction. The corrective gain led to a significant decrease in early stance vertical GRF by 2.3% and 4.4% of body weight for participants #2 and #3, respectively. Additionally, the late swing minimum knee flexion angles significantly increased by 2.78*deg* and 1.32*deg* for those two participants. These few-degree increases of minimum knee flexion help prevent any chance of hyperextension, while still avoiding the crouch-walking behavior that was observed using the Slow controller.

To summarize the effects of the nominal path speed, Γ, we have demonstrated a significant correlation between low nominal path speeds and unwanted behaviours such as crouch gait, as well as poor controller performance in terms of path tracking and excess energy absorption by the exoskeleton, which suggests disagreement between the intended movements of the user, and the exoskeleton. On the other hand, setting the nominal path speed too high, at the average path speed during the swing phase led to mild foot stomping and hyperextension. Our findings show that increasing the nominal path speed from its originally proposed value in [12] of Γ = 167, resulted in (1) enhanced comfort, natural behavior, and path speed, (2) lower path RMS error, (3) greater knee range of motion. We have demonstrated behaviours across a full spectrum of nominal path speeds and propose that the ideal nominal path speed should be time-varying according to the phase of the gait cycle, and 80% of the path speed observed during natural walking without an exoskeleton.

## IV. Conclusion

In this paper, we first analytically demonstrate that flow controller minimizes position and velocity kinematic errors. We introduced two modifications to flow controller based on participant’s (1) averaged or (2) instantaneous path speed, both of which improved controller performance by enabling more natural knee range of motion and decreasing the exoskeleton’s resistance to the user’s motions.

Experiments at three treadmill speeds showed that low nominal path speeds, including the originally proposed value [12], limit knee range of motion and can cause crouch gait. Increasing the nominal path speed (in Mean and Variable controllers) reduced path tracking error and increased knee range of motion, while also decreasing exoskeleton resistance to motion, as reflected in the power and energy profiles.

Participant feedback and quantitative analysis indicated that the modified controllers yielded more natural and comfortable gait than the original controller. Due to variability in individual gait patterns, particularly among motor-impaired users, reference path and path speed should be personalized. This includes adjusting the scaling of a corrective gain for Mean Γ as explored in this study. Inline with prior work [22], [23], we anticipate such personalization to effectively enhance the performance of the proposed modified flow controller. Future work will focus on implementing and testing personalized or adaptive flow controllers in individuals with motor impairments (e.g., those with incomplete spinal cord injury) to accommodate for their residual motor capacities and divergent gait patterns from able-bodied norms.

